## Supplementary File 1 for "Genomic epidemiology of syphilis reveals independent emergence of macrolide resistance across multiple circulating lineages"

R Notebook: Beale MA, et al. (2019) Genomic Epidemiology of Syphilis


Code 

- Show All Code
- Hide All Code
- Download Rmd

### R Notebook: Beale MA, et al. (2019) Genomic Epidemiology of Syphilis

### Import environment and library dependencies


```
R.Version()
```


```
$platform
[1] "x86_64-apple-darwin15.6.0"

$arch
[1] "x86_64"

$os
[1] "darwin15.6.0"

$system
[1] "x86_64, darwin15.6.0"

$status
[1] ""

$major
[1] "3"

$minor
[1] "5.1"

$year
[1] "2018"

$month
[1] "07"

$day
[1] "02"

$`svn rev`
[1] "74947"

$language
[1] "R"

$version.string
[1] "R version 3.5.1 (2018-07-02)"

$nickname
[1] "Feather Spray"
```


```
# import libraries
library(ggplot2)
library(ape)
#devtools::install_github("GuangchuangYu/ggtree")
library(ggtree)
library(phytools)
library(plyr)
library(dplyr)
library(reshape2)
library(grid)
library(gridExtra)
library(scales)
#library(phangorn)
library(randomcoloR)
#devtools::install_github("alexwailan/rpinecone")
library(rPinecone)
```


```
print(sessionInfo())
```


```
R version 3.5.1 (2018-07-02)
Platform: x86_64-apple-darwin15.6.0 (64-bit)
Running under: OS X El Capitan 10.11.6

Matrix products: default
BLAS: /System/Library/Frameworks/Accelerate.framework/Versions/A/Frameworks/vecLib.framework/Versions/A/libBLAS.dylib
LAPACK: /Library/Frameworks/R.framework/Versions/3.5/Resources/lib/libRlapack.dylib

locale:
[1] en_GB.UTF-8/en_GB.UTF-8/en_GB.UTF-8/C/en_GB.UTF-8/en_GB.UTF-8

attached base packages:
[1] grid      stats     graphics  grDevices utils     datasets  methods   base     

other attached packages:
 [1] bindrcpp_0.2.2    rPinecone_0.1.0   randomcoloR_1.1.0 scales_1.0.0      gridExtra_2.3    
 [6] reshape2_1.4.3    dplyr_0.7.8       plyr_1.8.4        phytools_0.6-60   maps_3.3.0       
[11] ggtree_1.14.6     ape_5.2           ggplot2_3.1.0    

loaded via a namespace (and not attached):
 [1] httr_1.4.0              tidyr_0.8.2             jsonlite_1.6            assertthat_0.2.0       
 [5] expm_0.999-3            rvcheck_0.1.3           animation_2.6           yaml_2.2.0             
 [9] progress_1.2.0          numDeriv_2016.8-1       pillar_1.3.1            lattice_0.20-38        
[13] glue_1.3.0              quadprog_1.5-5          phangorn_2.4.0          uuid_0.1-2             
[17] digest_0.6.18           RColorBrewer_1.1-2      colorspace_1.4-0        htmltools_0.3.6        
[21] Matrix_1.2-15           BMhyd_1.2-8             XML_3.98-1.16           pkgconfig_2.0.2        
[25] rncl_0.8.3              purrr_0.3.0             phylobase_0.8.4         corpcor_1.6.9          
[29] mvtnorm_1.0-8           tidytree_0.2.3          Rtsne_0.15              tibble_2.0.1           
[33] combinat_0.0-8          withr_2.1.2             lazyeval_0.2.1          mnormt_1.5-5           
[37] magrittr_1.5            crayon_1.3.4            evaluate_0.12           nlme_3.1-137           
[41] MASS_7.3-51.1           xml2_1.2.0              data.table_1.12.0       prettyunits_1.0.2      
[45] tools_3.5.1             TreeSim_2.3             hms_0.4.2               geiger_2.0.6.1         
[49] stringr_1.3.1           V8_1.5                  munsell_0.5.0           cluster_2.0.7-1        
[53] plotrix_3.7-4           ade4_1.7-13             compiler_3.5.1          clusterGeneration_1.3.4
[57] RNeXML_2.3.0            rlang_0.3.1             rstudioapi_0.9.0        subplex_1.5-4          
[61] igraph_1.2.2            base64enc_0.1-3         rmarkdown_1.11          labeling_0.3           
[65] gtable_0.2.0            deSolve_1.21            curl_3.3                R6_2.3.0               
[69] knitr_1.21              bindr_0.1.1             fastmatch_1.1-0         treeio_1.6.2           
[73] stringi_1.2.4           parallel_3.5.1          Rcpp_1.0.0              scatterplot3d_0.3-41   
[77] tidyselect_0.2.5        xfun_0.4                coda_0.19-2
```

### Specify data to use

`> ariba run databases/ARIBA/Treponema_23s__19-03-18/Treponema_23s.nichols.db2/ ./SRR3584965-b_1.fastq.gz ./SRR3584965-b_2.fastq.gz ./SRR3584965-b/`

`> ariba summary --cluster_cols assembled,ctg_cov --v_groups ariba_summary_known_vars *.report.tsv`


```
AMR.file <- "Source_Data_Treponema-Global_23s.summary.renamed.csv"
```


```
new.full.meta.file <- "Source_Data_Sample_meta_data3.tsv"
```


```
# Maximum likelihood tree (midpoint rooted) of all 137 genomes
iqtree.file <- "Source_Data_Treponema_Global_gubbins+iqtree_200718_SS14-fully-masked-.renamed.treefile"
```


```
# Beast maximum credibility tree for 110 clinically derived genomes
beast.file <- "Source_Data_Tpa-Global_SS14-mapped-gubbins_20072018.WGS.all-aln.Clinicals.renamed.noAr.no-invariant_StrictConstant_consensus.tree"
```


```
# joint ancestral reconstruction of SS14 sequences - snp scaled tree with midpoint rooting
pyjar.snpscale.tree.file <- "Source_Data_Tpa-Global_SS14-mapped-gubbins_20072018_Clinicals_pyjar.joint.renamed.tre"
```


```
# Penicillin binding protein gene variants source data (Ariba summary)
ariba.pen.file <- "Source_Data_ariba-penicillin.novel_21-11-2018.csv"
```


##### Filter reference mapped MSA for recombination using Gubbins

`> run_gubbins.py -c 4 --prefix Tpp-Global_SS14-mapped-gubbins_17072018 /lustre/scratch118/infgen/team216/mb29/Treponema/Treponema_Globals/gubbins/17-07-2018_SS14_full-masked/Treponema_pallidum_subsp_pallidum_SS14_v2_bwa.full-aln.fas`

##### Generate Maximum Likelihood phylogeny on Gubbins filtered SNP-alignment using IQ-Tree

`> iqtree -s /lustre/scratch118/infgen/team216/mb29/Treponema/Treponema_Globals/gubbins/20-02-18_SS14_fully-masked/Tpp-Global_SS14-mapped-gubbins_190218.filtered_polymorphic_sites.fasta -pre Treponema_Global_gubbins+iqtree_200218_map-to-SS14-fully-masked- -ntmax 4 -mem 8G -m MFP+MERGE+ASC -bb 1000`

#### Plot IQ-Tree (basic)


```
my.iqtree <- read.tree(iqtree.file)
my.iqtree <- midpoint.root(my.iqtree)
my.iqtree2 <- my.iqtree
# Clean up tip labels
my.iqtree2$tip.label <- gsub("\\|.+$","",gsub("TPP","",gsub("TPPCM\\_","",my.iqtree2$tip.label)))
# make basic
my.iqtree.plot <- ggtree(my.iqtree2)
my.iqtree.plot2 <- my.iqtree.plot +   
  # add in UF bootstrap support as node points
  geom_point2(aes(subset=(!isTip & as.numeric(label)>95))) + 
  # add tiplabels
  geom_tiplab(align=T,size=2,offset=0.001) + xlim_tree(0.1) + 
  geom_treescale(offset=-5,x=0.025,y=50)
```


```
Duplicated aesthetics after name standardisation: size
```


```
my.iqtree.plot2
```

#### Bring in AMR data and also metadata


```
full.meta <- read.csv(file=new.full.meta.file,comment.char = "", sep="\t", header=T)
colnames(full.meta)[1] <- "Sample"
full.meta.country <- full.meta[,c("Sample","Lineage","GeoCountry")]
colnames(full.meta.country) <- c("Sample","Lineage","Country")
# Clean up sample names
full.meta.country$Sample <- gsub("\\|.+$","",gsub("TPP","",gsub("TPPCM\\_","",full.meta.country$Sample)))
# make dataframe longform
full.meta.country.melt <- melt(full.meta.country,id.vars="Sample")
```


```
attributes are not identical across measure variables; they will be dropped
```


```
full.meta.country.melt[full.meta.country.melt$value=="-",] <- NA #### deal with missing data - for this plot will be empty
```

#### Now plot ML tree with metadata


```
# Define colors for metadata - can do this using distinctColorPalette - or specify a palette.
#countryLin.palette <- distinctColorPalette(length(unique(full.meta.country3.melt$value)),runTsne = F,altCol = F)
countryLin.palette <- c("#D8D9A3","#DD5691","#DE9260","#DADC5C","#80DEA2","royalblue2","#D7A3C4","indianred1","#7FE5DD","#BC79DA","#80A0E2")
# Make plot with metadata
my.iqtree.plot.heatmap <- facet_plot(my.iqtree.plot2, 'Meta', full.meta.country.melt, geom_tile, aes(x=as.numeric(as.factor(variable)),fill=as.factor(value)),width=0.75) +
  # specify colour pallete from earlier
  scale_fill_manual(breaks=unique(full.meta.country.melt$value),values=countryLin.palette) +
  theme(legend.position = "right") +
  theme(strip.background = element_rect(colour="white", fill="white"),strip.text.x = element_text(color="white")) +
  labs(fill="Country") +
  xlim_expand(10,'Meta') + 
  NULL
my.iqtree.plot.heatmap
```

### Now prepare to run rPinecone

Bring in data from precalculated SS14 tree (calculated in IQ-Tree), then analysed using joint ancestral reconstruction with pyjar. Branch lengths are scaled by SNPs, and the code also displays the number of SNPs on each branch


```
pyjar.snpscale.tree <- read.tree(pyjar.snpscale.tree.file)
# create a variable containing edge lengths and positions for plotting
edge <- data.frame(pyjar.snpscale.tree$edge, edge_num=1:length(pyjar.snpscale.tree$edge.length))
colnames(edge)=c("parent", "node", "edge_num")
edge$SNPs <- pyjar.snpscale.tree$edge.length
edge <- data.frame(pyjar.snpscale.tree$edge, edge_num=1:length(pyjar.snpscale.tree$edge.length))
colnames(edge)=c("parent", "node", "edge_num")
edge$SNPs <- pyjar.snpscale.tree$edge.length
# now build tree
pyjar.snpscale.tree.plot <- ggtree(pyjar.snpscale.tree) %<+% 
  edge + geom_text(aes(x=branch, label=SNPs, vjust=-.5),size=3,color="grey50") + 
  geom_tiplab(size=2,align=T,offset=.0001,linetype="dotted",linesize = 1) +
  #xlim(0, 80) +
  NULL
```


```
Duplicated aesthetics after name standardisation: size
```


```
pyjar.snpscale.tree.plot
```


#### Use rPinecone to determine SS14 clusters based on a root-to-tip defined threshold of 10 SNPs


```
# convert pyjar tree into a phylo object for reading into pinecone
pyjar.snpscale.tree.phylo <- as.phylo(pyjar.snpscale.tree)
# run pinecone
pinecone.output <- rPinecone::pinecone(pyjar.snpscale.tree.phylo,10,3) # same as 9,3
```


```
    104 taxa found from phylogenetic tree

Node_1  will be investigated.
Mexico_A|NAmerica|1953  skipped as this is a leaf.

Node_2  will be investigated.
CZ27|Europe|2012  skipped as this is a leaf.

Node_3  will be investigated.
Threhold met - Sub-lineage Number assigned:  1

Node_4  will be investigated.
TPPCM_UW370B|NAmerica|2005  skipped as this is a leaf.
TPPCM_UW187B|NAmerica|2003  skipped as this is a leaf.

Node_5  will be investigated.
TPPCM_UW213B|NAmerica|2003  skipped as this is a leaf.
TPPCM_UW254C|NAmerica|2004  skipped as this is a leaf.
TPPCM_UW231B|NAmerica|2004  skipped as this is a leaf.
TPPCM_UW116B|NAmerica|2002  skipped as this is a leaf.

Node_6  will be investigated.
TPPCM_UW254B|NAmerica|2004  skipped as this is a leaf.
TPPCM_UW215B|NAmerica|2004  skipped as this is a leaf.
TPPCM_UW228B|NAmerica|2004  skipped as this is a leaf.
TPPCM_UW186B|NAmerica|2003  skipped as this is a leaf.

Node_7  will be investigated.
TPPCM_UW330B|NAmerica|2005  skipped as this is a leaf.
TPPCM_UW337B|NAmerica|2005  skipped as this is a leaf.

Node_8  will be investigated.

Node_9  will be investigated.
SHD-R|Asia|2014  skipped as this is a leaf.
C3|Asia|2015  skipped as this is a leaf.
SHG-I2|Asia|2014  skipped as this is a leaf.
SHE-V|Asia|2014  skipped as this is a leaf.
Q3|Asia|2015  skipped as this is a leaf.

Node_10  will be investigated.
B3|Asia|2015  skipped as this is a leaf.
SHC-0|Asia|2014  skipped as this is a leaf.

Node_11  will be investigated.
Amoy|Asia|2011  skipped as this is a leaf.
K3|Asia|2015  skipped as this is a leaf.

Node_12  will be investigated.

Node_13  will be investigated.
Threhold met - Sub-lineage Number assigned:  2
TPPCM_UW281B|NAmerica|2004  skipped as this is a leaf.
PT_SIF1167|Europe|2013  skipped as this is a leaf.
TPPNL09|Europe|2016  skipped as this is a leaf.
PT_SIF1156|Europe|2013  skipped as this is a leaf.
TPPCM_UW140B|NAmerica|2002  skipped as this is a leaf.
PT_SIF0697|Europe|2009  skipped as this is a leaf.

Node_14  will be investigated.
Threhold met - Sub-lineage Number assigned:  3
TPPCM_UW368B|NAmerica|2005  skipped as this is a leaf.
TPPCM_UW149B|NAmerica|2003  skipped as this is a leaf.
TPPCM_UW125B|NAmerica|2002  skipped as this is a leaf.
TPPCM_UW181B|NAmerica|2003  skipped as this is a leaf.
TPPCM_UW298B|NAmerica|2004  skipped as this is a leaf.
TPPCM_UW138B|NAmerica|2002  skipped as this is a leaf.
TPPCM_UW262B|NAmerica|2004  skipped as this is a leaf.

Node_15  will be investigated.
TPPCM_UW134B|NAmerica|2002  skipped as this is a leaf.
TPPCM_UW104B|NAmerica|2002  skipped as this is a leaf.
TPPCM_UW099B|NAmerica|2001  skipped as this is a leaf.
TPPCM_UW133B|NAmerica|2002  skipped as this is a leaf.

Node_16  will be investigated.
TPPCM_UW257B|NAmerica|2004  skipped as this is a leaf.
TPPCM_UW344B|NAmerica|2005  skipped as this is a leaf.
TPPCM_UW126B|NAmerica|2002  skipped as this is a leaf.

Node_17  will be investigated.
TPPCM_UW147B|NAmerica|2003  skipped as this is a leaf.
TPPCM_UW102B|NAmerica|2002  skipped as this is a leaf.

Node_18  will be investigated.
Threhold met - Sub-lineage Number assigned:  4
TPPCM_UW093B|NAmerica|2001  skipped as this is a leaf.
TPPCM_UW155B|NAmerica|2003  skipped as this is a leaf.
TPPCM_UW211B|NAmerica|2003  skipped as this is a leaf.

Node_19  will be investigated.

Node_20  will be investigated.
Threhold met - Sub-lineage Number assigned:  5

Node_21  will be investigated.
PT_SIF0857|Europe|2010  skipped as this is a leaf.
PT_SIF1196|Europe|2013  skipped as this is a leaf.
PT_SIF0908|Europe|2010  skipped as this is a leaf.

Node_22  will be investigated.
PT_SIF1242|Europe|2014  skipped as this is a leaf.
PT_SIF1002|Europe|2011  skipped as this is a leaf.

Node_23  will be investigated.
TPPCM_UW291B|NAmerica|2004  skipped as this is a leaf.

Node_24  will be investigated.
TPPCM_UW195B|NAmerica|2003  skipped as this is a leaf.

Node_25  will be investigated.
TPPCM_UW280B|NAmerica|2004  skipped as this is a leaf.
TPPCM_UW244B|NAmerica|2004  skipped as this is a leaf.
TPPCM_UW376B|NAmerica|2005  skipped as this is a leaf.

Node_26  will be investigated.
TPPCM_UW391B|NAmerica|2005  skipped as this is a leaf.
TPPCM_UW391C|NAmerica|2005  skipped as this is a leaf.

Node_27  will be investigated.
Threhold met - Sub-lineage Number assigned:  6
PT_SIF1063|Europe|2012  skipped as this is a leaf.

Node_28  will be investigated.
PT_SIF1020|Europe|2011  skipped as this is a leaf.
PT_SIF1135|Europe|2013  skipped as this is a leaf.
NE19|Europe|2013  skipped as this is a leaf.

Node_29  will be investigated.
Threhold met - Sub-lineage Number assigned:  7
TPPCM_UW248B|NAmerica|2004  skipped as this is a leaf.
TPPCM_UW492B|NAmerica|2006  skipped as this is a leaf.

Node_30  will be investigated.
TPPNL16|Europe|2016  skipped as this is a leaf.
SW6|Europe|2012  skipped as this is a leaf.

Node_31  will be investigated.
TPPCM_UW148B|NAmerica|2003  skipped as this is a leaf.
TPPCM_UW473B|NAmerica|2006  skipped as this is a leaf.

Node_32  will be investigated.
Threhold met - Sub-lineage Number assigned:  8
PT_SIF1183|Europe|2013  skipped as this is a leaf.
TPPCM_UW074B|NAmerica|2004  skipped as this is a leaf.
TPPCM_UW259B|NAmerica|2004  skipped as this is a leaf.
TPPCM_UW411B|NAmerica|2006  skipped as this is a leaf.
PT_SIF0751|Europe|2009  skipped as this is a leaf.
NE17|Europe|2013  skipped as this is a leaf.
PT_SIF1140|Europe|2013  skipped as this is a leaf.
PT_SIF1280|Europe|2014  skipped as this is a leaf.
TPPNL10|Europe|2016  skipped as this is a leaf.

Node_33  will be investigated.
TPPCM_UW327B|NAmerica|2005  skipped as this is a leaf.
TPPCM_UW264B|NAmerica|2004  skipped as this is a leaf.

Node_34  will be investigated.
NE15|Europe|2013  skipped as this is a leaf.
TPPNL11|Europe|2016  skipped as this is a leaf.

Node_35  will be investigated.
PT_SIF0877_3|Europe|2010  skipped as this is a leaf.
PT_SIF1261|Europe|2014  skipped as this is a leaf.

Node_36  will be investigated.
PT_SIF1142|Europe|2013  skipped as this is a leaf.
PT_SIF1299|Europe|2014  skipped as this is a leaf.

Node_37  will be investigated.
PT_SIF1278|Europe|2014  skipped as this is a leaf.
PT_SIF1252|Europe|2014  skipped as this is a leaf.

Node_38  will be investigated.
TPPNL13|Europe|2016  skipped as this is a leaf.

Node_39  will be investigated.
SW8|Europe|2012  skipped as this is a leaf.
AU16|Europe|2013  skipped as this is a leaf.
AU15|Europe|2013  skipped as this is a leaf.

Node_40  will be investigated.
TPPCM_UW397B|NAmerica|2006  skipped as this is a leaf.
TPPCM_UW304B|NAmerica|2005  skipped as this is a leaf.
TPPCM_UW303B|NAmerica|2005  skipped as this is a leaf.
TPPCM_UW379B|NAmerica|2005  skipped as this is a leaf.
TPPCM_UW823B|NAmerica|2011  skipped as this is a leaf.
PT_SIF1200|Europe|2013  skipped as this is a leaf.

Node_41  will be investigated.
TPPCM_UW383B|NAmerica|2005  skipped as this is a leaf.
PT_SIF0954|Europe|2010  skipped as this is a leaf.

Node_42  will be investigated.
TPPCM_UW526B|NAmerica|2007  skipped as this is a leaf.
TPPCM_UW852B|NAmerica|2011  skipped as this is a leaf.
TPPCM_UW824B|NAmerica|2011  skipped as this is a leaf.

Number of Isolates on tree:  104
Number of Sub-lineages identified:  8
Number of Major Sublineages identified:  1
Number of Singletons remain:  2
Major Sub-Lineages  1 is composed of  7  Sub-lineages &  81  isolates.
```


rPinecone groups the SS14 sequences into 1 major sublineages and 8 minor sublineages. We use the minor sublineages.


```
# Take key pinecone lineage data from a list into a dataframe
pinecone.clusters <- data.frame(pinecone.output$table[,c(1,2)],stringsAsFactors = T)
# harmonise tip names
tiplab.names <- data.frame(pyjar.snpscale.tree$tip.label)
colnames(tiplab.names) <- "Taxa"
pinecone.clusters<- plyr::join(pinecone.clusters,tiplab.names)
```


```
Joining by: Taxa
```


```
# create dataframe for plotting
rownames(pinecone.clusters) <- pinecone.output$table[,1]
colnames(pinecone.clusters) <- c("Sample","Sublineage")
pinecone.clusters$Sublineage <- as.factor(pinecone.clusters$Sublineage)
# pinecone labels each singleton as an individual (e.g. 'singleton_1', 'singleton_2') - want to deprecate this behaviour here
pinecone.clusters$Sublineage <- gsub("singleton.+$","singleton",pinecone.clusters$Sublineage)
```


rPinecone represents the tree structure reasonably well, but for clarity, we wished to break lineage 1 (which is geographically distinct, as well as distinct based on different resistance alleles). We therefore subclassified samples in sublineage 1 into 1A or 1B based on ancestral node


```
# Further manually subdivide lineage 1 based on ancestral nodes 109 and 120
pinecone.clusters$Sublineage <- sapply(1:nrow(pinecone.clusters), function(x) ifelse(pinecone.clusters$Sublineage[x]==1 & pyjar.snpscale.tree.plot$data$node[x] %in% unlist(phangorn::Descendants(as.phylo(pyjar.snpscale.tree.plot),109,"tips")),"1A",as.character(pinecone.clusters$Sublineage[x])))
pinecone.clusters$Sublineage <- sapply(1:nrow(pinecone.clusters), function(x) ifelse(pinecone.clusters$Sublineage[x]==1 & pyjar.snpscale.tree.plot$data$node[x] %in% unlist(phangorn::Descendants(as.phylo(pyjar.snpscale.tree.plot),120,"tips")),"1B",as.character(pinecone.clusters$Sublineage[x])))
# prepare new dataframe for plotting
pinecone.clusters$Sublineage <- as.factor(pinecone.clusters$Sublineage)
pinecone.clusters2 <- data.frame(pinecone.clusters[,2])
rownames(pinecone.clusters2) <-  as.character(pinecone.clusters$Sample)
colnames(pinecone.clusters2) <- "Sublineage"
```


Rough plot of pinecone clusters vs pyjar tree


```
# plot tree with metadata (rough)
pyjar.snpscale.tree.plot %>% gheatmap(pinecone.clusters2,offset=15,colnames_position='top',colnames_offset_y=2,font.size=2,width=0.035)
```

### Plot basic BEAST tree


```
# Bring in beast tree and extract tree data into dataframe
my.beast.tree <- read.beast(beast.file)
my.beast.tree.data <- fortify(my.beast.tree)
# Plot minimally annotated BEAST Tree (no tip labels or colours)
beast.tree.plot3 <- ggtree(my.beast.tree,mrsd="2016-06-01",ladderize = T) + 
  theme_tree2() +
# Add date lines for easy interpretation  
  scale_x_continuous(breaks=c(1700,1750, 1900, 1930,1960,1980, 2000, 2015), minor_breaks=seq(1960, 2016, 5)) +
  theme(panel.grid.major   = element_line(color="grey50", size=.2),
        panel.grid.minor   = element_line(color="grey85", size=.2),
        panel.grid.major.y = element_blank(),
        panel.grid.minor.y = element_blank()) #+ xlim_tree(2090)
# Add posterior support as node points
beast.tree.plot3 <- beast.tree.plot3 + geom_point2(aes(subset=(!isTip & as.numeric(posterior)>0.8)),color="gray60",size=3,alpha=0.5) + 
  geom_point2(aes(subset=(!isTip & as.numeric(posterior)>0.91)),color="gray40",size=3,alpha=0.5) + 
  geom_point2(aes(subset=(!isTip & as.numeric(posterior)>=0.96)),color="black",size=3,alpha=0.5)
#beast.tree.plot3 <- rotate(beast.tree.plot3,117)
beast.tree.plot3
```


```
# Get dates for divergence of SS14 and Nichols
node.mrca <- 110 # specify node in tree
# Mean date
2016.5 - my.beast.tree.data[my.beast.tree.data$node==110,"height"]
```


```
    height
1 1749.732
```


```
# Median date 
2016.5 - my.beast.tree.data[my.beast.tree.data$node==110,"height_median"]
```


```
  height_median
1      1755.304
```


```
# 95% HPD (confidence intervals)
2016.5 - as.numeric(unlist(my.beast.tree.data[my.beast.tree.data$node==110,"height_0.95_HPD"]))
```


```
[1] 1835.444 1651.452
```


#### Now bring in macrolide data and combine with lineage info from pinecone and tree


```
# Read in macrolide resistance analysis from ARIBA
my.AMR <- read.csv(AMR.file,header=T)
# Need to filter duplicates in the table
my.AMR <- my.AMR[!duplicated(my.AMR$name),]
# Or filter those that are missing from the tree
sample.labels <- data.frame(my.iqtree$tip.label)
colnames(sample.labels) <- "fullname"
my.AMR2 <- merge(sample.labels,my.AMR, by.x="fullname", by.y="name")
my.AMR2 <- my.AMR2[!duplicated(my.AMR2$fullname),]
## 
rownames(my.AMR2) <- my.AMR2$fullname
# Deal with ARIBA entries without frequency scores (i.e. negatives which are blanked to 'NA')
my.AMR2$NR_076156.23S.2058G.. <- sapply(1:nrow(my.AMR2), function (x) ifelse(my.AMR2$NR_076156.23S.2058G[x]=="no" & is.na(my.AMR2$NR_076156.23S.2058G..[x]),0,my.AMR2$NR_076156.23S.2058G..[x]))
my.AMR2$NR_076156.23S.2059G.. <- sapply(1:nrow(my.AMR2), function (x) ifelse(my.AMR2$NR_076156.23S.2059G[x]=="no" & is.na(my.AMR2$NR_076156.23S.2059G..[x]),0,my.AMR2$NR_076156.23S.2059G..[x]))
# Clean up headings
colnames(my.AMR2) <- gsub("\\.\\.","%",gsub("NR\\_076156\\.","",colnames(my.AMR2)))
my.AMR2 <- my.AMR2[,c("fullname","23S.2058G%","23S.2059G%")]
colnames(my.AMR2) <- c("Sample","A2058G","A2059G")
# Ariba data frequency outputs can be used quantitatively, but for these purposes, simplify variant frequency to sensitive, mixed or resistant (and if a variant is <5% or >95%, treat it as 0/100)
my.AMR2.cat <- my.AMR2
my.AMR2.cat$A2058G <- ifelse(my.AMR2.cat$A2058G<5,0,ifelse(my.AMR2.cat$A2058G>=95,100,50))
my.AMR2.cat$A2059G <- ifelse(my.AMR2.cat$A2059G<5,0,ifelse(my.AMR2.cat$A2059G>=95,100,50))
# merge back together
my.AMR2.cat.pinecone <- plyr::join(my.AMR2.cat,pinecone.clusters,type='full')
```


```
Joining by: Sample
```


```
my.AMR2.melt <- melt(my.AMR2.cat,id.vars="Sample")
# define colours for resistance columns
fine = 3
macro.palette = colorRampPalette(c("white","gray60","black"))
my.AMR2.melt$mycolor = macro.palette(fine)[as.numeric(cut(my.AMR2.melt$value,breaks = fine))]
my.AMR2.melt <- my.AMR2.melt[order(my.AMR2.melt$value),]
# Clean up pinecone values - singleton samples that don't form part of a sublineage will plot as white
pinecone.clusters.melt <- melt(pinecone.clusters,id.vars="Sample")
pinecone.clusters.melt[pinecone.clusters.melt$value=="singleton","value"] <- NA
pinecone.clusters.melt$mycolor <- distinctColorPalette(length(unique(pinecone.clusters.melt$value)),runTsne = F,altCol = F)[as.numeric(cut(as.numeric(pinecone.clusters.melt$value),breaks =length(unique(pinecone.clusters.melt$value))-1))]
```


```
NAs introduced by coercion
```


```
AMR.pinecone <- rbind(pinecone.clusters.melt,my.AMR2.melt)
full.meta2 <- full.meta
full.meta2$SampleShort <- full.meta2$Sample
full.meta2$Sample <- full.meta2$TipName
# clean up dataframes
full.meta.AMR.pincone <- plyr::join(my.AMR2.cat.pinecone, full.meta2)
```


```
Joining by: Sample
```


```
my.AMR2.cat.pinecone2 <- full.meta.AMR.pincone[,c("Sample","Lineage","Sublineage","A2058G","A2059G")]
my.AMR2.cat.pinecone2.melt <- melt(my.AMR2.cat.pinecone2,id.vars="Sample")
```


```
attributes are not identical across measure variables; they will be dropped
```


```
subL.AMR.cols <- c("white","black","#DE9260","#C3E3A5","#A3F5CC","#FAEC80","#7AA1E5","#C31AB9","#999999","#23A73E","#B56C7D","#FED1AA","royalblue2","white","indianred1")
```

#### Now do BEAST + metadata plot


```
facet1 <- facet_plot(beast.tree.plot3, 'heatmap', my.AMR2.cat.pinecone2.melt[!grepl("A205",my.AMR2.cat.pinecone2.melt$variable),], geom_tile,aes(x=as.numeric(as.factor(variable)),fill=as.factor(value),color=NULL),width=0.75) 
facet2 <- facet_plot(facet1, 'heatmap', my.AMR2.cat.pinecone2.melt[grepl("A205",my.AMR2.cat.pinecone2.melt$variable),], geom_tile,aes(x=as.numeric(as.factor(variable)),fill=as.factor(value)),color="grey10",width=0.75)
facet.done <- facet2 + scale_fill_manual(breaks=c(unique(AMR.pinecone$value)),values=c(subL.AMR.cols)) + 
  theme(legend.position = "right") +
  theme(strip.background = element_rect(colour="white", fill="white"),strip.text.x = element_text(color="white")) +
  labs(fill="Meta") +
  xlim_expand(7.5,'heatmap') +
  NULL
facet.done
```

#### Plot clade showing sublineages 1A and 1B


```
#Plot subclade
subL.AMR.cols.subtree <- subL.AMR.cols[c(1:12,14)]
my.AMR2.cat.pinecone3 <- my.AMR2.cat.pinecone[,c("Sublineage","A2058G","A2059G")]
rownames(my.AMR2.cat.pinecone3) <- my.AMR2.cat.pinecone$Sample
#113, 195
viewClade(beast.tree.plot3, node=197) %>% gheatmap(my.AMR2.cat.pinecone3,width=0.025,color='grey45',font.size = 4.5,hjust=0.8,colnames_position='top',colnames_offset_y=1.5,colnames_angle=-45) + 
  scale_fill_manual(breaks=c(unique(AMR.pinecone$value)),values=c(subL.AMR.cols.subtree))
```


```
attributes are not identical across measure variables;
they will be droppedScale for 'fill' is already present. Adding another scale for 'fill', which will replace the
existing scale.
```


```
#facet.done + geom_text2(aes(subset=!isTip, label=node), hjust=-.3)
```

### Now look at data by pinecone clusters


```
###########
# Do some statistics using pinecone clusters
meta.pinecone.binary <- full.meta.AMR.pincone
meta.pinecone.binary$A2058G <- ifelse(meta.pinecone.binary$A2058G>=50,1,0)
meta.pinecone.binary$A2059G <- ifelse(meta.pinecone.binary$A2059G>=50,1,0)
meta.pinecone.binary$Resistant <- ifelse((meta.pinecone.binary$A2058G==1 |meta.pinecone.binary$A2059G==1) ,1,0)
meta.pinecone.binary$Year <- as.numeric(gsub("^.+\\|","",meta.pinecone.binary$Sample))
```


```
NAs introduced by coercion
```


```
## Circle plot
meta.pinecone.binary.summ <- meta.pinecone.binary %>% dplyr::group_by(Sublineage,Year) %>% dplyr::count(Sublineage,Year)
meta.pinecone.binary.summ <- meta.pinecone.binary.summ[!is.na(meta.pinecone.binary.summ$Sublineage),]
p.circle <- ggplot(meta.pinecone.binary.summ, aes(x=Year, y=Sublineage,size=n)) + geom_point(alpha=1/2, aes(color=Sublineage)) + theme_minimal() + xlim(1980,2020) 
p.circle
```


#### Extract sublineage ancestral nodes and plot (in red)


```
# Manually assign ancestral nodes for pinecone lineages using tree generated with above code (pinecone doesn't provide this yet)
# Redefined after tree revision 07-2018
pinecone.mrca.nodes <- data.frame(
         Sublineage = c("1A","1B","2","3","4","5","6","7","8"),
      mrca.node = c(207,199, 175, 182, 180, 128, 139, 122, 142)
   )
# Show node labels and selected node points on beast tree 
facet.done + 
  geom_text2(aes(subset=!isTip, label=node), hjust=-.3) +
  geom_point2(aes(subset=(node %in% pinecone.mrca.nodes$mrca.node)),color="red") +
  ggtitle("Red nodes indicate MRCA for each sublineage")
```


```
#Extract relevant nodes data from beast tree data
pinecone.mrca.nodes.beast <- merge(pinecone.mrca.nodes,my.beast.tree.data[,c("node","height","height_0.95_HPD","height_median","height_range")], by.x="mrca.node",by.y="node")
# Data is in the form of "height" information - need to convert to years relative to mrcd (2016/06/01)
pinecone.mrca.nodes.beast$mrca.median <- 2016.5 - pinecone.mrca.nodes.beast$height_median
pinecone.mrca.nodes.beast$year <- as.numeric(round(2016.5 - pinecone.mrca.nodes.beast$height_median,0))
pinecone.mrca.nodes.beast$mrca.95high <- round(2016.5 - sapply(1:nrow(pinecone.mrca.nodes.beast),function(x) as.numeric(unlist(pinecone.mrca.nodes.beast[x,"height_0.95_HPD"]))[1]))
pinecone.mrca.nodes.beast$mrca.95low <- round(2016.5 - sapply(1:nrow(pinecone.mrca.nodes.beast),function(x) as.numeric(unlist(pinecone.mrca.nodes.beast[x,"height_0.95_HPD"]))[2]))
# Color Sublineages using same scheme as above
sublin.col <- subL.AMR.cols[c(3,4,5,6,7,8,10,11,12)]
```


```
# Show dates for lineages
pinecone.mrca.nodes.beast[order(pinecone.mrca.nodes.beast$Sublineage),c("Sublineage","mrca.node","year","mrca.95high","mrca.95low")]
```


```
  Sublineage mrca.node year mrca.95high mrca.95low
9         1A       207 1990        1997       1981
8         1B       199 2002        2008       1993
5          2       175 2001        2002       1996
7          3       182 1987        1995       1976
6          4       180 1999        2001       1995
2          5       128 1996        2001       1990
3          6       139 2006        2010       2000
1          7       122 2000        2003       1995
4          8       142 1996        2001       1990
```


```
###
# Need to factor and reorder Sublineages to ensure they display in vertically descending order
meta.pinecone.binary.summ <- meta.pinecone.binary.summ[!is.na(meta.pinecone.binary.summ$Sublineage),]
colnames(meta.pinecone.binary.summ)[3] <- "Count"
meta.pinecone.binary.summ$Sublineage <- factor(meta.pinecone.binary.summ$Sublineage,levels=rev(sort(unique(meta.pinecone.binary.summ$Sublineage))))
pinecone.mrca.nodes.beast$Sublineage <- factor(pinecone.mrca.nodes.beast$Sublineage,levels=rev(sort(unique(pinecone.mrca.nodes.beast$Sublineage))))
pinecone.mrca.nodes.beast$yaxis <- unique(pinecone.mrca.nodes.beast$Sublineage)
```

#### Now plot all sublineages with MRCA (and confidence intervals)


```
p.circle <- ggplot() + 
  # First plot the sample dates
  geom_point(data=meta.pinecone.binary.summ[meta.pinecone.binary.summ$Sublineage!="singleton",], aes(x=Year, y=Sublineage,size=Count,color=Sublineage),alpha=0.5) + 
  # Colour the samples using the same scheme as for the BEAST tree
  scale_colour_manual(values = rev(sublin.col)) +
  # Then plot the TMRCA
  geom_point(data=pinecone.mrca.nodes.beast, aes(x=year, y=Sublineage),fill="black",color="black",size=3) + 
  # Finally plot the confidence intervals
  geom_rect(data=pinecone.mrca.nodes.beast,aes(xmin=mrca.95low, xmax=mrca.95high, ymin=as.numeric(Sublineage)-0.1, ymax=as.numeric(Sublineage)+0.1,group=Sublineage),alpha=0.25,fill="red") + 
  # And clean up the labels and theme
  labs(x="Year",y="Sublineage") +
  theme_bw() + coord_cartesian(xlim=c(1975,2020))
p.circle
```

#### Now look at resistance by sublineage


```
my.AMR2.cat.pinecone3 <- my.AMR2.cat.pinecone2[!is.na(my.AMR2.cat.pinecone2$Sublineage),]
my.AMR2.cat.pinecone3$Resistant <- ifelse((my.AMR2.cat.pinecone3$A2058G==100 | my.AMR2.cat.pinecone3$A2059G==100) ,"Resistant",ifelse((my.AMR2.cat.pinecone3$A2058G==50 | my.AMR2.cat.pinecone3$A2059G==50),"Mixed","Sensitive"))
my.AMR2.cat.pinecone3$Sublineage <- factor(my.AMR2.cat.pinecone3$Sublineage,levels=rev(sort(unique(my.AMR2.cat.pinecone3$Sublineage))))
p.pinecone.resistance <- ggplot(my.AMR2.cat.pinecone3[my.AMR2.cat.pinecone3$Sublineage!="singleton",], aes(Sublineage,group=Resistant,fill=factor(Resistant))) + geom_bar(stat='count',position="fill",color="black",width=0.5) + 
  theme_bw() + 
  scale_fill_manual(breaks=c("Sensitive","Mixed","Resistant"),values=c("gray60","black","white")) + 
  labs(y="Proportion",x="Sublineage", fill="A2058G/A2059G")
p.pinecone.resistance + coord_flip()
```


Alternative way of plotting resistance (break up by resistance allele)


```
my.AMR2.cat.pinecone4 <- my.AMR2.cat.pinecone2[!is.na(my.AMR2.cat.pinecone2$Sublineage),]
my.AMR2.cat.pinecone4$Resistant <- ifelse((my.AMR2.cat.pinecone4$A2058G==100),"A2058G",
                                          ifelse((my.AMR2.cat.pinecone4$A2059G==100),"A2059G",ifelse((my.AMR2.cat.pinecone4$A2058G==50 | my.AMR2.cat.pinecone4$A2059G==50),"Mixed","Sensitive")))
my.AMR2.cat.pinecone4$Sublineage <- factor(my.AMR2.cat.pinecone4$Sublineage,levels=rev(sort(unique(my.AMR2.cat.pinecone4$Sublineage))))
p.pinecone.resistance2 <- ggplot(my.AMR2.cat.pinecone4[my.AMR2.cat.pinecone4$Sublineage!="singleton",], aes(Sublineage,group=Resistant,fill=factor(Resistant))) + geom_bar(stat='count',position="fill",color="black",width=0.5) + 
  theme_bw() + 
  scale_fill_manual(breaks=c("Sensitive","Mixed","A2058G","A2059G"),values=c("black","gray30","gray70","white")) + 
  labs(y="Count",x="Sublineage", fill="A2058G/A2059G")
p.pinecone.resistance2 + coord_flip()
```


##### Construct plot

Final editing of legends was performed in Inkscape


```
grid.arrange(p.circle + theme(legend.position = "left"), p.pinecone.resistance+coord_flip() + theme(legend.position = "none", axis.title.y = element_blank(), axis.text.y = element_blank(), axis.ticks.y= element_blank()) ,nrow=1, widths=c(3,1))
```

#### Extract useful stats


```
# Final dataset for stats (only include those in the trees)
full.meta.AMR.pincone.inTree <- full.meta.AMR.pincone[!is.na(full.meta.AMR.pincone$TipName),]
full.meta.AMR.pincone.inTree <- plyr::rename(full.meta.AMR.pincone.inTree,c("Recent_Clinical"="Clinical"))
```


```
# Total sequences in total dataset (IQ-Tree)
nrow(full.meta.AMR.pincone.inTree)
```


```
[1] 122
```


```
# Total sequences from this study and published elsewhere
nrow(full.meta.AMR.pincone.inTree[full.meta.AMR.pincone.inTree$Type=="WSI",])
```


```
[1] 73
```


```
nrow(full.meta.AMR.pincone.inTree[full.meta.AMR.pincone.inTree$Type=="Public",])
```


```
[1] 49
```


```
# Number of UK and North American samples sequenced in this study
nrow(full.meta.AMR.pincone.inTree[(full.meta.AMR.pincone.inTree$Type=="WSI" & full.meta.AMR.pincone.inTree$GeoCountry=="UK"),])
```


```
[1] 8
```


```
nrow(full.meta.AMR.pincone.inTree[(full.meta.AMR.pincone.inTree$Type=="WSI" & full.meta.AMR.pincone.inTree$GeoCountry=="USA"),])
```


```
[1] 65
```


```
# Number of USA Samples that are also Clinical
nrow(full.meta.AMR.pincone.inTree[(full.meta.AMR.pincone.inTree$Type=="WSI" & full.meta.AMR.pincone.inTree$GeoCountry=="USA" & full.meta.AMR.pincone.inTree$Clinical=="Yes"),])
```


```
[1] 60
```


```
# Note that 5 non-clinical samples sequenced in this study had been previously published elsewhere. Consensus sequences were near identical (small differences in Seattle_81-4), and phylogenetic placement was also the same. Since some of these genomes have no associated citation, and to allow the authors freedom to publish unhindered, we chose to use our version of these genomes in all analyses.   
as.character(full.meta.AMR.pincone.inTree[full.meta.AMR.pincone.inTree$Type=="WSI" & full.meta.AMR.pincone.inTree$Clinical=="No","SampleShort"])
```


```
[1] "Chicago_tube2"     "Nichols_Houston_E" "Nichols_Houston_J" "Nichols_Houston_O" "Seattle_81-4"
```


```
# Number of clinical samples by sequencing source
nrow(full.meta.AMR.pincone.inTree[full.meta.AMR.pincone.inTree$Type=="WSI" & full.meta.AMR.pincone.inTree$Clinical=="Yes",])
```


```
[1] 68
```


```
nrow(full.meta.AMR.pincone.inTree[full.meta.AMR.pincone.inTree$Type=="Public" & full.meta.AMR.pincone.inTree$Clinical=="Yes",])
```


```
[1] 41
```


```
# Total Recent Clinical sequences in total dataset (and in BEAST tree)
nrow(full.meta.AMR.pincone.inTree[full.meta.AMR.pincone.inTree$Clinical=="Yes",])
```


```
[1] 109
```


```
# SS14 sequences in the total dataset (IQ-Tree)
nrow(full.meta.AMR.pincone.inTree[full.meta.AMR.pincone.inTree$Lineage=="SS14",])
```


```
[1] 105
```


```
# SS14 sequences that are clinical (in the BEAST tree)
nrow(full.meta.AMR.pincone.inTree[(full.meta.AMR.pincone.inTree$Lineage=="SS14"&full.meta.AMR.pincone.inTree$Clinical=="Yes"),])
```


```
[1] 103
```


```
# Nichols sequences in the total dataset (IQ-Tree)
nrow(full.meta.AMR.pincone.inTree[full.meta.AMR.pincone.inTree$Lineage=="Nichols",])
```


```
[1] 17
```


```
# Nichols sequences that are clinical (in the BEAST tree)
nrow(full.meta.AMR.pincone.inTree[(full.meta.AMR.pincone.inTree$Lineage=="Nichols"&full.meta.AMR.pincone.inTree$Clinical=="Yes"),])
```


```
[1] 6
```


```
# Number of samples from USA
nrow(full.meta.AMR.pincone.inTree[(full.meta.AMR.pincone.inTree$GeoCountry=="USA"),])
```


```
[1] 72
```


```
# Number of samples from UK
nrow(full.meta.AMR.pincone.inTree[(full.meta.AMR.pincone.inTree$GeoCountry=="UK"),])
```


```
[1] 8
```


```
# Number of samples from China
nrow(full.meta.AMR.pincone.inTree[(full.meta.AMR.pincone.inTree$GeoCountry=="China"),])
```


```
[1] 9
```


```
# Number of samples from Portugal
nrow(full.meta.AMR.pincone.inTree[(full.meta.AMR.pincone.inTree$GeoCountry=="Portugal"),])
```


```
[1] 23
```


```
# Date range
as.character(sort(unique(full.meta.AMR.pincone.inTree$Sample_Date)))
```


```
 [1] "-"    "1912" "1953" "1973" "1977" "1980" "2001" "2002" "2003" "2004" "2005" "2006" "2007" "2009"
[15] "2010" "2011" "2012" "2013" "2014" "2015" "2016"
```

```
# Number of resistant samples
full.meta.AMR.pincone.inTree$resistant <- ifelse((as.numeric(full.meta.AMR.pincone.inTree$A2058G)==0 & as.numeric(full.meta.AMR.pincone.inTree$A2059G)==0),"Sensitive",ifelse((as.numeric(full.meta.AMR.pincone.inTree$A2058G)==50 | as.numeric(full.meta.AMR.pincone.inTree$A2059G)==50),"Mixed","Resistant"))
#
nrow(full.meta.AMR.pincone.inTree[full.meta.AMR.pincone.inTree$resistant=="Resistant",])
```


```
[1] 76
```


```
nrow(full.meta.AMR.pincone.inTree[full.meta.AMR.pincone.inTree$resistant=="Mixed",])
```


```
[1] 7
```


```
nrow(full.meta.AMR.pincone.inTree[full.meta.AMR.pincone.inTree$resistant=="Sensitive",])
```


```
[1] 39
```


```
# Percentage resistant (all)
percent(nrow(full.meta.AMR.pincone.inTree[full.meta.AMR.pincone.inTree$resistant=="Resistant",])/nrow(full.meta.AMR.pincone.inTree))
```


```
[1] "62.3%"
```


```
#
# Percentage resistant (Clinical)
percent(nrow(full.meta.AMR.pincone.inTree[(full.meta.AMR.pincone.inTree$resistant=="Resistant" & full.meta.AMR.pincone.inTree$Clinical=="Yes") ,])/nrow(full.meta.AMR.pincone.inTree[full.meta.AMR.pincone.inTree$Clinical=="Yes",]))
```


```
[1] "68.8%"
```


```
# Number of resistant Nichols
nrow(full.meta.AMR.pincone.inTree[(full.meta.AMR.pincone.inTree$resistant=="Resistant" & full.meta.AMR.pincone.inTree$Lineage=="Nichols"),])
```


```
[1] 6
```


```
#
# % of Resistant Nichols
percent(nrow(full.meta.AMR.pincone.inTree[(full.meta.AMR.pincone.inTree$resistant=="Resistant" & full.meta.AMR.pincone.inTree$Lineage=="Nichols"),]) / nrow(full.meta.AMR.pincone.inTree[(full.meta.AMR.pincone.inTree$Lineage=="Nichols"),]))
```


```
[1] "35.3%"
```


```
# Number of Resistant SS14
nrow(full.meta.AMR.pincone.inTree[(full.meta.AMR.pincone.inTree$resistant=="Resistant" & full.meta.AMR.pincone.inTree$Lineage=="SS14"),])
```


```
[1] 70
```


```
#
# % of Resistant SS14
percent(nrow(full.meta.AMR.pincone.inTree[(full.meta.AMR.pincone.inTree$resistant=="Resistant" & full.meta.AMR.pincone.inTree$Lineage=="SS14"),]) / nrow(full.meta.AMR.pincone.inTree[(full.meta.AMR.pincone.inTree$Lineage=="SS14"),]))
```


```
[1] "66.7%"
```


```
# % of Resistant Nichols (Clinical only)
percent(nrow(full.meta.AMR.pincone.inTree[(full.meta.AMR.pincone.inTree$resistant=="Resistant" & full.meta.AMR.pincone.inTree$Lineage=="Nichols" & full.meta.AMR.pincone.inTree$Clinical=="Yes"),]) / nrow(full.meta.AMR.pincone.inTree[(full.meta.AMR.pincone.inTree$Lineage=="Nichols"& full.meta.AMR.pincone.inTree$Clinical=="Yes"),]))
```


```
[1] "100%"
```


```
# % of Resistant SS14 (Clinical only)
percent(nrow(full.meta.AMR.pincone.inTree[(full.meta.AMR.pincone.inTree$resistant=="Resistant" & full.meta.AMR.pincone.inTree$Lineage=="SS14" & full.meta.AMR.pincone.inTree$Clinical=="Yes"),]) / nrow(full.meta.AMR.pincone.inTree[(full.meta.AMR.pincone.inTree$Lineage=="SS14"& full.meta.AMR.pincone.inTree$Clinical=="Yes"),]))
```


```
[1] "67.0%"
```


```
# Number of Resistant SS14 with A2058G
nrow(full.meta.AMR.pincone.inTree[(full.meta.AMR.pincone.inTree$A2058G==100 & full.meta.AMR.pincone.inTree$Lineage=="SS14"),])
```


```
[1] 59
```


```
# Number of Resistant SS14 with A2059G
nrow(full.meta.AMR.pincone.inTree[(full.meta.AMR.pincone.inTree$A2059G==100 & full.meta.AMR.pincone.inTree$Lineage=="SS14"),])
```


```
[1] 11
```


```
# Look at samples with mixed resistance alleles
as.character(full.meta.AMR.pincone.inTree[full.meta.AMR.pincone.inTree$resistant=="Mixed","SampleShort"])
```


```
[1] "PT_SIF0857"   "PT_SIF0877_3" "PT_SIF1140"   "PT_SIF1142"   "PT_SIF1261"   "PT_SIF1278"  
[7] "NL13"
```


Write data out to table


```
write.table(full.meta.AMR.pincone.inTree,file="Full.metadata+AMR.csv",sep=",",row.names=F,quote=T)
```

### Perform additional analysis to look at penicillin binding protein variants

Run ARIBA to look for novel variants in pbp1 (TPANIC\_0500), mrcA (TPANIC\_0705 - mrcA), pbp2 (TPANIC\_0760), tp47 (TPANIC\_0574)

Example: `ariba run pen.genes.ariba-db/ Treponema_Globals/fastqs/22931_6#13.ds2405106-reads_1.fastq.gz Treponema_Globals/fastqs/22931_6#13.ds2405106-reads_2.fastq.gz Treponema_Globals/ARIBA/manual_penicillins/22931_6#13.ds2405106-reads`

Collate: `ariba summary --novel_variants --no_tree --cluster_cols assembled,novel_var,ctg_cov,pct_id,ref_seq ariba-penicillin.novel_21-11-2018 *.ariba.tsv`


```
ariba.pen <- read.csv(ariba.pen.file,sep=",",header=T)
# Fix gene names (each is labelled as clusterx in input data)
colnames(ariba.pen) <- gsub("cluster\\.","TPANIC_0705.",gsub("cluster\\_3","TPANIC_0574",gsub("cluster\\_2","TPANIC_0500",gsub("cluster\\_1","TPANIC_0760",colnames(ariba.pen)))))
# merge with existing metadata
ariba.pen.merge <- merge(full.meta2, ariba.pen, by.x="Subsampled_fastq",by.y="name")
# subset pen data to clinical samples in tree only
ariba.pen.merge.beasttree <- merge(ariba.pen.merge,data.frame(label=my.beast.tree.data[my.beast.tree.data$isTip==T,"label"],stringsAsFactors = F), by.x="Sample", by.y="label")
# Remove some of the unnecessary columns
ariba.pen.merge.beasttree.vars1 <- ariba.pen.merge.beasttree[,!grepl("ref\\_seq",colnames(ariba.pen.merge.beasttree))]
ariba.pen.merge.beasttree.vars1 <- ariba.pen.merge.beasttree.vars1[,!grepl("pct\\_id",colnames(ariba.pen.merge.beasttree.vars1))]
ariba.pen.merge.beasttree.vars1 <- ariba.pen.merge.beasttree.vars1[,!grepl("ctg\\_cov",colnames(ariba.pen.merge.beasttree.vars1))]
ariba.pen.merge.beasttree.vars1 <- ariba.pen.merge.beasttree.vars1[,!grepl("novel\\_var",colnames(ariba.pen.merge.beasttree.vars1))]
ariba.pen.merge.beasttree.vars1 <- ariba.pen.merge.beasttree.vars1[,!grepl("assembled",colnames(ariba.pen.merge.beasttree.vars1))]
ariba.pen.merge.beasttree.vars1 <- ariba.pen.merge.beasttree.vars1[,c(grepl("TPANIC",colnames(ariba.pen.merge.beasttree.vars1)))]
# Remove putative variant sites that don't occur in the clinical samples in the tree
variable.sites <- sapply(1:ncol(ariba.pen.merge.beasttree.vars1), function (y) length(unique(as.character(unlist(ariba.pen.merge.beasttree.vars1[,y])))))
ariba.pen.merge.beasttree.vars1.var <- ariba.pen.merge.beasttree.vars1[,variable.sites==2]
row.names(ariba.pen.merge.beasttree.vars1.var) <- ariba.pen.merge.beasttree$Sample
# reorder colnames according to gene and gene position
test.genes <- c("TPANIC_0500","TPANIC_0574","TPANIC_0705","TPANIC_0760")
all.vars <- NULL
for (current.gene in test.genes){
  current.var <- data.frame(var=colnames(ariba.pen.merge.beasttree.vars1.var)[grepl(current.gene,colnames(ariba.pen.merge.beasttree.vars1.var))],stringsAsFactors = F)
  current.var$pos <- as.numeric(gsub("[A-Z]+$","",gsub("^.+\\.[A-Z]{1}","",current.var$var, perl=T)))
  current.var <- current.var[order(current.var$pos),]
  all.vars <- rbind(all.vars, current.var)
}
ariba.pen.merge.beasttree.vars1.var <- ariba.pen.merge.beasttree.vars1.var[,all.vars$var]
colnames(ariba.pen.merge.beasttree.vars1.var) <- gsub("TPANIC","TP",colnames(ariba.pen.merge.beasttree.vars1.var))
# Plot tree with variants
beast.tree.plot3 %>% gheatmap(ariba.pen.merge.beasttree.vars1.var,
                              colnames_position='top',colnames_angle = -90,colnames_offset_y=8,font.size=2) +
  NULL
```

### Perform additional analysis to show transcontinental admixture amongst SS14 sublineages


```
# reorder lineages for plot
full.meta.AMR.pincone.inTree.2 <- full.meta.AMR.pincone.inTree
full.meta.AMR.pincone.inTree.2$Sublineage <- factor(full.meta.AMR.pincone.inTree.2$Sublineage,levels=rev(sort(unique(full.meta.AMR.pincone.inTree.2$Sublineage))))
# make plot of continent proportions by sublineage
p.pinecone.continent <- ggplot(full.meta.AMR.pincone.inTree.2[(full.meta.AMR.pincone.inTree.2$Sublineage!="singleton" & !is.na(full.meta.AMR.pincone.inTree.2$Sublineage)),], aes(Sublineage,group=Continent,fill=factor(Continent))) + geom_bar(stat='count',position="fill",color="black",width=0.5) + 
  theme_bw() + 
  labs(y="Proportion",x="Sublineage", fill="Continent")
p.pinecone.continent + coord_flip()
```

### Add resistance plot


```
grid.arrange(
  p.pinecone.resistance + coord_flip() + theme(legend.position = "left") + labs(y="Proportion") + ggtitle("Resistance Genotype"), 
             p.pinecone.continent + coord_flip() + theme(legend.position = "right", axis.title.y = element_blank(), axis.text.y = element_blank(), axis.ticks.y= element_blank()) + ggtitle("Geographical Origin"),
  ncol=2)
```

### Add penicillin data to metadata file and print to file


```
ariba.pen.merge.beasttree.vars1.var.2 <- ariba.pen.merge.beasttree.vars1.var
ariba.pen.merge.beasttree.vars1.var.2$Sample <- row.names(ariba.pen.merge.beasttree.vars1.var.2)
full.meta.AMR.pincone.inTree.pen <- plyr::join(full.meta.AMR.pincone.inTree,ariba.pen.merge.beasttree.vars1.var.2,by="Sample")
write.table(full.meta.AMR.pincone.inTree.pen,file="Full.metadata+AMR+pen.csv",sep=",",row.names=F,quote=T)
#full.meta.AMR.pincone.inTree.pen
```
